# Effects of conceptus proteins on endometrium and blood leukocytes of dairy cattle using transcriptome and meta-analysis

**DOI:** 10.1101/2024.04.25.591148

**Authors:** Maria Isabel da Silva, Troy Ott

**Affiliations:** Department of Animal Science, Center for Reproductive Biology and Health, Huck Institutes of the Life Sciences, Pennsylvania State University, University Park, PA, USA

**Keywords:** IFNT, PAG, genes, endometrium, blood, leukocytes, pregnancy, bovine

## Abstract

This study investigates the short and long-term effects of IFNT and PAG on the transcriptome of endometrium and blood leukocytes. Holstein heifers received intrauterine infusions of one of the following treatments: 20 mL of a 200 μg/mL bovine serum albumin solution (BSA; vehicle) from day 14 to 16 of the estrous cycle (BSA), vehicle + 10 μg/mL of IFNT from day 14 to 16 (IFNT3), vehicle + 10 μg/mL of IFNT from day 14 to 19 (IFNT6), and vehicle + 10 μg/mL of IFNT from day 14 to 16 followed by vehicle + 10 μg/mL of IFNT + 5 μg/mL of PAG from day 17 to 19 (IFNT+PAG). RNA-seq analysis was performed in endometrial biopsies and blood leukocytes collected after treatments. Acute IFNT signaling in the endometrium (IFNT3 vs BSA), induced differentially expressed genes (DEG) associated with interferon activation, immune response, inflammation, cell death, and inhibited vesicle transport and extracellular matrix remodeling. Prolonged IFNT signaling (IFNT6 vs IFNT3) altered gene expression related to cell invasion, retinoic acid signaling, and embryo implantation. In contrast, PAG induced numerous DEG in blood leukocytes but only 4 DEG in the endometrium. In blood leukocytes, PAG stimulated genes involved in development and TGFB signaling while inhibiting interferon signaling and cell migration. Overall, IFNT is a primary regulator of endometrial gene expression, while PAG predominantly affected the transcriptome of circulating immune cells during early pregnancy. Further research is essential to fully grasp the roles of identified DEG in both the endometrium and blood leukocytes.

**Graphic abstract:** 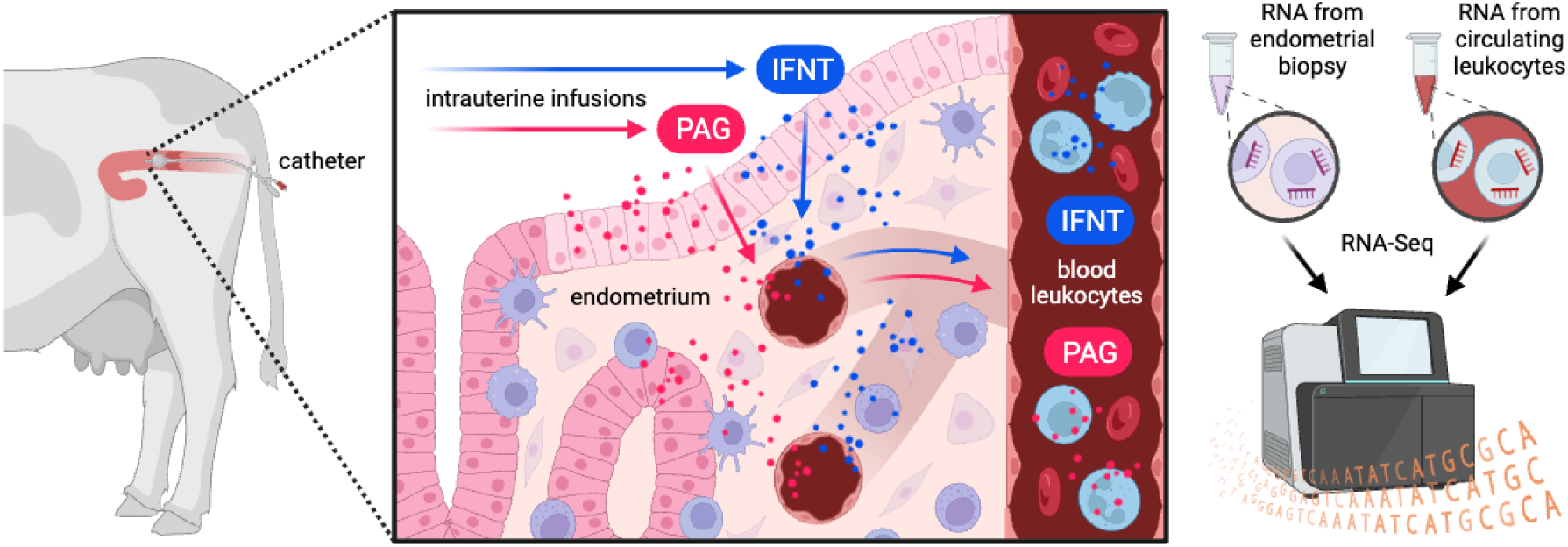

**Main findings:**

- Acute IFNT signaling in the endometrium activates genes involved in inflammation while inhibiting vesicle transport in cells and ECM remodeling.
- Prolonged IFNT signaling in the endometrium regulates genes involved in cell invasion and retinoic acid signaling.
- PAG alter gene expression in blood leukocytes more than in endometrium and may stimulate leukocyte differentiation while inhibiting leukocyte extravasation.

## INTRODUCTION

The first 4 weeks of pregnancy are a decisive period for embryo survival (Diskin et al., 2016). The day 6-8 blastocyst depends on endometrial secretions stimulated by progesterone (P4) to elongate into a filamentous conceptus by day 16 (Brenner & West, 1975; Brooks et al., 2014; Flechon et al., 1986). Throughout elongation and up to day 38 of pregnancy, the conceptus secretes interferon tau (IFNT) that mediates maternal recognition of pregnancy by altering endometrial synthesis and release of prostaglandin F2 alpha (PGF2A) to maintain the corpus luteum (CL) and P4 secretion (Bazer et al., 1997; Bazer & Thatcher, 2017; McCracken et al., 1970). The conceptus begins to attach to the endometrium around day 18-21 via interdigitation of microvilli on the chorion and endometrial epithelium and then multinucleated trophoblast cells migrate and fuse with the luminal epithelium and later establish placentomes where the chorion is in contact with endometrial caruncles (King et al., 1980; Wimsatt, 1951; Wooding, 1992; Wooding & Burton, 2008). As trophoblast cells fuse with epithelial cells, they release granules containing pregnancy associated glycoproteins (PAG) into the endometrial stroma (Wooding, 1992). The role of PAG is still unknown, but they increase in concentration in circulation throughout pregnancy (Wallace et al., 2015). Concurrently, the conceptus must also evade immune attack by altering immune cell phenotype to create an immunotolerant uterine environment and promote uterine remodeling to form the placenta (Ott, 2019). Thus, pregnancy is established through highly orchestrated conceptus-maternal interactions and immune adaptation.

Interferon tau is critical for CL maintenance which is accomplished by inhibiting transcription of endometrial estrogen receptor alpha (ESR1) necessary for the pulsatile release of luteolytic PGF2A (Spencer et al., 1995). Previous transcriptome studies showed that interferon tau increases the expression of endometrial genes involved in activation of an antiviral response, interferon gamma (IFNG) signaling, ion transport, and apoptosis (Bauersachs et al., 2012; Chaney et al., 2021; Forde et al., 2011; Forde et al., 2012; Mathew et al., 2019). Compared to interferon stimulated genes (ISG), interferon downregulated genes (IDG) in the endometrium were fewer in number and related to inhibition of Notch signaling, angiogenesis, corticotrophin releasing hormone signaling, cell migration, WNT/B-catenin signaling, phagocytosis, synapsis and cAMP response element (CREB) signaling in the endometrium (Bauersachs et al., 2012; Chaney et al., 2021; Forde et al., 2011; Forde et al., 2012; Mathew et al., 2019). However, these studies have limitations that may conceal important effects of IFNT on endometrial function. Some limitations include use of interferon alpha (IFNA), *in vitro* culture, and/or a short time treatment that do not reflect the physiology of IFNT secretion in pregnancy. Another limitation is the use of microarray assays that profile predefined genes via hybridization.

The time of appearance of PAG in the endometrium also suggests that these proteins may play a role during conceptus attachment and placentation. The role of PAG is still unknown, but some evidence suggests that they may regulate immune function, cell adhesion, and extracellular matrix (ECM) remodeling (Wallace et al., 2015, 2019). To date, only one transcriptome study described gene expression around the time of the initiation of PAG secretion. It was observed that, on day 20 of pregnancy compared to day 20 of estrous cycle, pathways related to interferon signaling, antigen presentation, leukocyte extravasation, and tight junctions were enriched in caruncular endometrium (CAR) while pathways for protein ubiquitination, oxidative phosphorylation, antigen presentation, and vascular endothelial growth factor (VEGF) signaling were enriched in intercaruncular endometrium (ICAR) (Mansouri-Attia et al., 2009). However, the design in this study did not allow determination of a causative role for PAG in these pathways. We conducted an experiment to determine the effects of IFNT and PAG on gene expression in the endometrium and peripheral blood leukocytes. Treatments were planned to evaluate the effects of a short- and longer-term IFNT exposure as well as the additive effect of PAG along IFNT exposure. We hypothesized that, while short-term IFNT signaling activates immune response pathways, prolonged IFNT signaling may alter gene expression to facilitate implantation and, in the presence of PAG, may activate transcription of additional genes associated with placentation and uterine remodeling. In blood leukocytes, we hypothesized that PAG inhibit immune cell proliferation and migration.

## MATERIALS AND METHODS

### Ethics statement

This research was conducted with cattle from the Dairy Barn of Pennsylvania State University. Experimental procedures and animal handling were conducted in accordance with the Guide for the Care and Use of Agricultural Animals in Agricultural Research and Teaching and approved by the Pennsylvania State University.

### The experiment

Postpubertal Holstein heifers (N=30; 12-17 months of age) were estrous synchronized with an intramuscular injection of a PGF2A analog (Estroplan, 500 μg of cloprostenol sodium, Parnell) and were monitored for estrus behavior (day 0). Thirteen days after the onset of estrus, heifers were moved to tie-stall pens and fed a typical heifer total mixed ration once daily. In the morning of day 14, animals received caudal epidural anesthesia using lidocaine (5-8 mL of 2% lidocaine hydrochloride, Aspen Veterinary Resources) and their perineal area was cleaned. A sterilized cervical dilator (5 mm shaft diameter, Minitube GmbH) coated with Chlorhexidine lube (VetOne) was passed through the cervix. Then, a 16 Fr silicone Foley catheter (Bardex) mounted over a sterilized artificial insemination pipette plunger was inserted through the cervix and secured in the uterine body via inflation of the catheter balloon with 7 mL of sterile phosphate-buffered saline (PBS; pH 7.7) solution. The insemination pipette plunger was removed and a sterilized tubing connector with custom-made cap was placed into the drainage channel to seal catheter from environmental contamination. Transrectal ultrasound was performed to confirm catheter placement.

Vehicle solution was prepared with 200 μg/mL bovine serum albumin (BSA) fraction V (OmniPur) in PBS, sterilized using a vacuum filtration system with polyethersulfone (PES) membrane (2 μm pore size; VWR), and stored at 4°C. Each treatment infusion had a total volume of 20 mL and consisted of either vehicle only, vehicle + 200 μg of IFNT (10 μg/mL), or vehicle + 200 μg of IFNT + 100 μg of PAG (5 μg/mL). Treatments were prepared using recombinant ovine IFNT (kindly supplied by Fuller Bazer, Texas A&M University) and PAG purified from first-trimester bovine placenta (kindly supplied by Josh Branen, Biotracking LLC Inc). The amount of IFNT infused was determined based on previous literature reporting IFNT production in 24 hours culture of day 16 ovine embryos (Ashworth and Bazer, 1989). Amount of PAG infused was determined via ELISA assay (BioPRYN, Biotracking LLC Inc, Moscow, Idaho) using uterine flushes of heifers on day 20 of pregnancy (N=4, ∼3 μg/mL of PAG). Treatments were freshly prepared daily and brought to room temperature prior to infusion.

Heifers (N=6 per treatment) were randomly assigned to one of the following treatment schedules: vehicle from day 14 to 16 of the estrous cycle (BSA), vehicle + IFNT from day 14 to 16 (IFNT3), vehicle + IFNT from day 14 to 19 (IFNT6), and vehicle + IFNT from day 14 to 16 followed by vehicle + IFNT + PAG from day 17 to 19 (IFNT+PAG). Before application of each infusion, the capped end of catheters was cleaned with 70% ethanol. Intrauterine treatments were applied twice daily at 12-hour intervals (8:30 am and 8:30 pm). After each infusion, 5 mL of sterile PBS solution was infused though catheter to ensure that full treatment volume was injected in the uterus. Animals were inspected daily for signs of uterine infection such as vaginal discharge and elevated rectal temperatures.

Prior to morning infusions, blood was collected daily into K3 EDTA vacuette tubes (Greiner Bio-one) for plasma analysis and isolation of peripheral blood leukocytes (PBL) as described in Kamat et al., (2016). At the end of the treatment schedule, 12 hours after the last infusion (8:30 am of day 17 or day 20), blood was also collected as described above. Then, animals received caudal epidural anesthesia and had their perineal area cleaned. A sterilized custom-made mini-Tischler biopsy forceps with a 22 inch shaft length, 4 mm shaft diameter, and 4x8 mm bite size (Aries surgical, Davis CA) was passed through the cervix for collection of endometrial biopsies (N=2). Biopsies were immediately placed into sterilized cryotubes, snap-frozen in liquid nitrogen, and stored at -80°C.

Animals were removed from the study if the catheter became dislodged during treatment schedule (N=2) or if they exhibited purulent vaginal discharge on the day of biopsy (N=3). At the end of experiment, all animals received an injection of PGF2A analog to induce estrus and minimize uterine infections. They were also monitored for 3 days after the experiment and received veterinary care when showing abnormal health (N=1). Of the 24 heifers that successfully completed the experiment, 75% (N=18) became pregnant within 2 services after returning to the herd.

### Progesterone and PAG ELISA

Plasma progesterone concentration was measured in all animals to determine the presence and/or maintenance of a functional of corpus luteum during the experimental period. Plasma was isolated by centrifuging 9 mL of blood at 1,500 x g for 15 minutes and progesterone concentration was determined using an ELISA protocol and reagents previously validated for cattle (Hughes et al., 2021). Plasma PAG concentration was measured from day 17 to 20 in animals that received IFNT+PAG to assess movement of PAG across the uterine wall and into the circulation as occurs during pregnancy. PAG concentration was also measured in stock plasma of heifers (N=6) and cows (N=5) on days 17 and 20 of pregnancy from a previous experiment to compare appearance and concentrations of PAG in circulation between treated heifers and pregnant heifers and cows. All pregnant animals delivered live and healthy calves. Plasma samples were shipped in dry ice to Biotracking LLC Inc and PAG were measured using a commercially available sandwich ELISA assay (BioPRYN, Biotracking LLC Inc, Moscow, Idaho).

### RNA isolation and sequencing

RNA was isolated from one frozen endometrial biopsy (∼50-60 mg) and PBL (15 million cells) following homogenization in 800 µL of TRIzol reagent (Life Technologies). Total RNA was purified using RNA Clean and Concentrator kit (Zymo Research) to eliminate contaminating DNA and reagents. Endometrial RNA samples from all treatment groups (N=4 per treatment) and PBL RNA samples from IFNT6 and IFNT+PAG (N=4 per treatment) were shipped on dry ice to BGI laboratory (Beijing Genomics Institute, Shenzhen, China) where RNA quality and quantity were assessed using an Agilent 2100 Bioanalyzer (Agilent Technologies). All samples met the quality criteria for RNA sequencing, showing RNA integrity number (RIN) between 8 and 9.9 values. Preparation of mRNA library and sequencing were performed following BGI DNBSEQ technology platform with paired end 100 bp (PE100).

### RNA-seq analysis

Transcriptome data were processed according to BGI bioinformatic pipeline. Raw reads were filtered using SOAPnuke v1.5.2 software (Cock et al., 2010) and samples averaged 92% clean reads. Clean reads were aligned to the reference genome GCF_000003205.7_Btau_5.0.1 (*Bos taurus*, NCBI) using HISAT2 v2.0.4 software (D. Kim et al., 2015) for identification of novel transcripts with an average alignment ratio of 90%. Clean reads were mapped to the annotated transcriptome using Bowtie2 v2.2.5 software (Langmead & Salzberg, 2012) with an average alignment ratio of 70%. A total of 18,801 genes were detected. Then, gene expression of each sample was calculated from reads mapped to the transcriptome using RSEM v1.2.8 software (B. Li & Dewey, 2011), and the DEseq2 method (Love et al., 2014) was used for detection of differentially expressed genes (DEG) in the following comparison groups: (1) IFNT3 versus BSA, (2) IFNT6 versus IFNT3, and (3) IFNT+PAG versus IFNT6. For the IFNT3 versus BSA comparison, DEG were considered significant if Log2 FC ≥ |1| (same as ≥ |2| fold change) and Q-value of < 0.05. However, for the IFNT6 versus IFNT3 and IFNT+PAG versus IFNT6 comparisons, DEG were considered significant if Log2 FC ≥ |1| and Q-value < 0.15 due to small differences in the composition of compared treatments, variability between replicates and the desire to minimize the probability of Type II errors.

### Data analysis

Functional analysis was performed in Ingenuity Pathway Analysis (IPA; Qiagen) using the DEG from IFNT3 vs BSA and IFNT6 vs IFNT3 comparisons. This analysis was used to identify canonical pathways, upstream regulators, and causal networks related to early embryo loss that were affected by treatments. Results from IPA analysis was considered significant if Fisher’s exact test had a p-value < 0.05 and status prediction had a z-score ≥ |2|. Moreover, upstream regulators that were present in the DEG list were selected if the directionality of expression (upregulation or downregulation) corresponded with predicted status (activated or inhibited). Heatmaps of top, abundant, upregulated and downregulated DEG were created using the Morpheus software (https://software.broadinstitute.org/morpheus) for matrix visualization and analysis.

The list of DEG in IFNT3 was uploaded to the INTERFEROME database (http://www.interferome.org/interferome/home.jspx) and compared with the list of genes known to be regulated by other interferons. The search was conducted in November 2022 with comparison filters set to include all studies with any type of interferon, under any experimental conditions, and a minimum fold change of |1|. Additionally, DEG from our results were compared with DEG from other studies using a free Venn diagram tool (https://bioinformatics.psb.ugent.be/webtools/Venn/). Studies were selected if they evaluated the endometrial gene expression profile during bovine early pregnancy via RNA-seq and/or RNA-microarray analysis. The generated list of common and/or unique genes across studies were further analyzed using IPA comparison analysis.

### Validation of DEG via RT-qPCR

Reverse transcription quantitative polymerase chain reaction (RT-qPCR) was conducted with selected DEG from IFNT+PAG vs IFNT6 comparison. This analysis was done using endometrial and PBL RNA from all animals (N=6/treatment) treated with IFNT+PAG and IFNT6 as well as endometrial RNA (N=12/group) and PBL RNA (N=7/group) from day 17 cyclic (D17C), day 17 pregnant (D17P), and day 20 pregnant (D20P) heifers from previous experiments. Primers (table) were designed using the National Center for Biotechnology Information (NCBI) primer blast (https://www.ncbi.nlm.nih.gov/tools/primer-blast/) and validated via Sanger sequencing of amplicons (Penn State University, Huck Institutes of the Life Sciences Genomics Core Facility). Ribosomal protein L19 (RPL19) and actin beta (ACTB) were selected as reference genes based on their abundant and stable expression in endometrium and PBL. All RNA samples were treated with DNase using the RQ1 RNase-free DNase kit (Promega). Synthesis of cDNA was performed using the AzuraQuant cDNA synthesis kit (Azura Genomics), and real-time polymerase chain reaction (RT-qPCR) was conducted using AzuraQuant Green Fast qPCR Mix LoRox (Azura Genomics), according to manufacturer’s protocols. Signal intensity was captured using QuantStudio 3 qPCR system (Applied Biosystems) under the following conditions: (1) initial denaturation at 95°C for 2 minutes, (2) 40 cycles of 95°C for 5 seconds, 60°C for 30 seconds, 72°C for 30 seconds, and (3) final extension at 72°C for 2 minutes. For each primer set, samples were run in duplicate along a standard curve consisting of 8 log dilutions of the PCR product to confirm assay performance and detection limit. Standard curves showed efficiencies between 88% to 100%, and samples with cycle threshold below detection limit of the standard curve were removed from statistical analysis. Statistical analyses were performed with 2^−ΔΔCt^ (fold change) values using an unpaired Student’s t-test or a one-way ANOVA. Least square means (LSM) ± standard error of means (SEM) were graphed using GraphPad 9.3 (GraphPad Software Inc.).

## RESULTS

### Plasma analysis across experimental replicates

All treated and control animals had plasma progesterone concentrations that were greater than 4 ng/mL (Figure 2.1A). This confirmed that intrauterine infusion of recombinant ovine IFNT inhibited luteolysis, especially in heifers from IFNT6 and IFNT+PAG. For the vehicle control treatment group, as expected, progesterone concentrations had not started to decline on the morning of day 17 when the endometrium was biopsied. Detection of PAG in the plasma was observed in 3/6 heifers from IFNT+PAG at a concentration of ∼ 200 pg/mL after day 17 (Figure 2.1B). In comparison, pregnant heifers did not have detectable PAG in plasma on day 17, but on day 20 of pregnancy PAG were present in plasma of 3/6 heifers at a concentration of ∼ 140 pg/mL (Figure 2.1C). All pregnant cows had circulating PAG detected on day 17 and 20, both at a concentration of ∼ 340 pg/mL (Figure 2.1D). Overall, the concentrations of PAG in plasma were similar between treated and pregnant heifers. Interestingly, our results suggest that cows may have earlier and/or greater secretion of PAG than heifers. However, this hypothesis requires future investigation as it cannot be ruled out that the higher concentrations of PAG at day 17 in cows resulted from residual PAG from a prior pregnancy.

**Figure 1:**
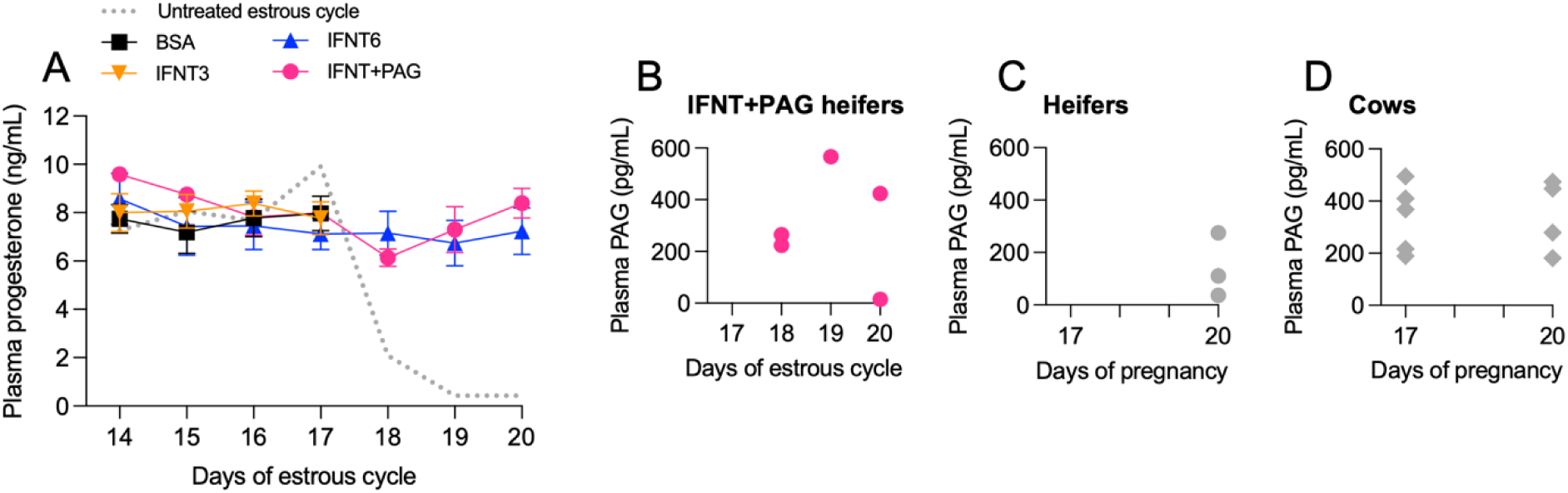
Plasma progesterone and PAG concentrations across experimental replicates. (A) Mean concentration of P4 in plasma of heifers from all treatment groups (N=6/treatment) and untreated estrous cycle (N=1). (B) Mean concentration of PAGs in plasma of IFNT+PAG heifers, (C) pregnant heifers, and (D) pregnant cows.

**Figure 2:**
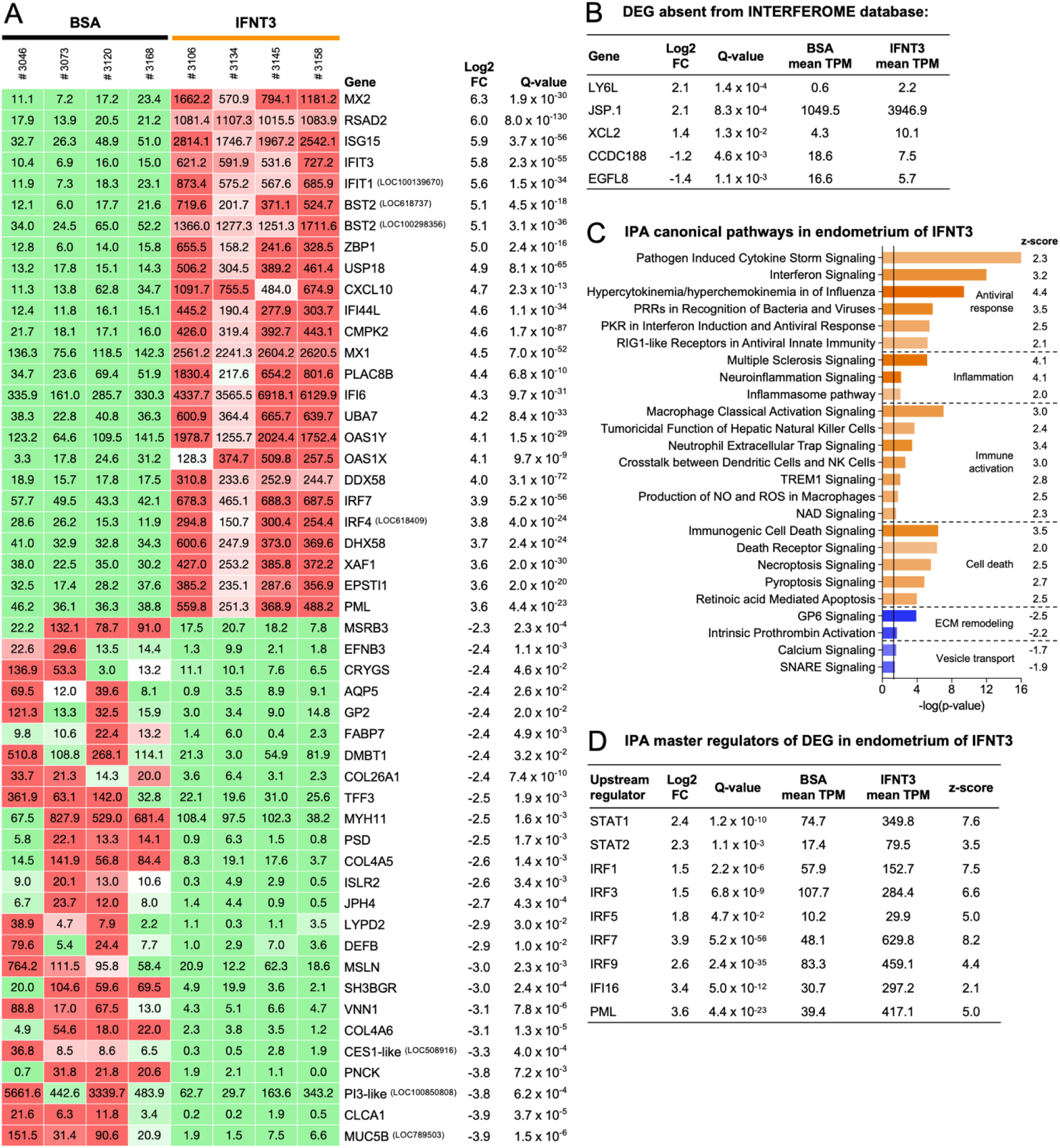
Differentially expressed genes (DEG), canonical pathways and upstream regulators identified in endometrium of IFNT3 vs BSA. (A) Upregulated and downregulated in DEG with transcripts per million (TPM) values in each animal replicate (#) shown inside the rectangles. (B) DEG not found in the INTERFEROME database that may be specifically regulated by IFNT in the bovine endometrium. (C) Main IPA canonical pathways predicted to be activated (positive z-score, orange) and inhibited (negative z-score, blue) by IFNT treatment with significance threshold -log (p-value of 0.05) = 1.3 (solid line). (D) Main IPA upstream regulators of transcription predicted to be activated in response to IFNT treatment.

### Endometrial DEG in IFNT3 vs BSA

There were a total of 1,101 DEG in IFNT3 compared with BSA. The 602 upregulated DEG consisted of 506 characterized transcripts, 84 predicted transcripts (LOC), 7 uncharacterized transcripts, and 5 transcripts homologous to chromosome regions in the bovine genome. Similarly, 499 downregulated DEG consisted of 463 characterized transcripts, 21 predicted transcripts (LOC), 10 uncharacterized transcripts, and 5 transcripts homologous to chromosome regions in the bovine genome. The top 20 up and down DEG with the greatest number of TPM are shown in Figure 2A. Using the INTERFEROME database, all characterized DEG in our dataset have been identified as type I, II and/or III interferon-regulated genes, except for LY6L, JSP.1, XCL2, CCDCC118, and EGFL8 (Figure 2B). These 5 genes may represent novel interferon regulated genes specific to our experimental conditions or simply not be directly regulated by interferons.

The IPA database recognized 489 upregulated and 465 downregulated DEG from our dataset. These DEG overlapped with a total of 21 activated and 4 inhibited IPA-based canonical pathways (p-value < 0.05, z-score ≥ |1.7|) indicating that, overall, IFNT3 induced activation of interferon and viral response, inflammation, immune activation, and cell death while it inhibited extracellular matrix (ECM) remodeling and vesicle transport in the endometrium (Figure 2C). Upstream analysis identified 9 activated transcription factors (p-value < 0.05, z-score ≥ 2) that were both DEG and could directly regulate other genes from our dataset in response to IFNT3 (Figure 2D). IPA causal network analysis predicted that IFNT3 treatment inhibited nuclear receptor corepressor 1 (NCOR1) as a master regulator of DEG in IFNT3 (p-value < 0.05, z-score ≤ -2). However, NCOR1 transcript was not detected in our analysis.

Additionally, when comparing our results with experimentally similar studies (Figure 3A) we observed 96 upregulated DEG common to our dataset and those studies (Figure 3B) and only 23 downregulated DEG common to at least one similar study (Figure 3C). Note that differences in DEG across studies may result from not only discrepancies in experimental design, but also differences in methods (RNA-microarray vs DNA-seq), gene nomenclature and discovery at the time each study was conducted. Due to the low number of downregulated genes across published studies, further comparisons were done using only upregulated DEG to identify genes regulated by the conceptus/pregnancy, type-I interferon, other interferons, IFNT in CAR and ICAR in epithelial cells (EC), stromal fibroblast (SF) and other cell types (Figure 2.3D). Overall, independent of experimental conditions, IFNT was predicted to regulate functions of immune cells including migration, cell death, proliferation, and phenotype polarization (Figure 3E).

**Figure 3:**
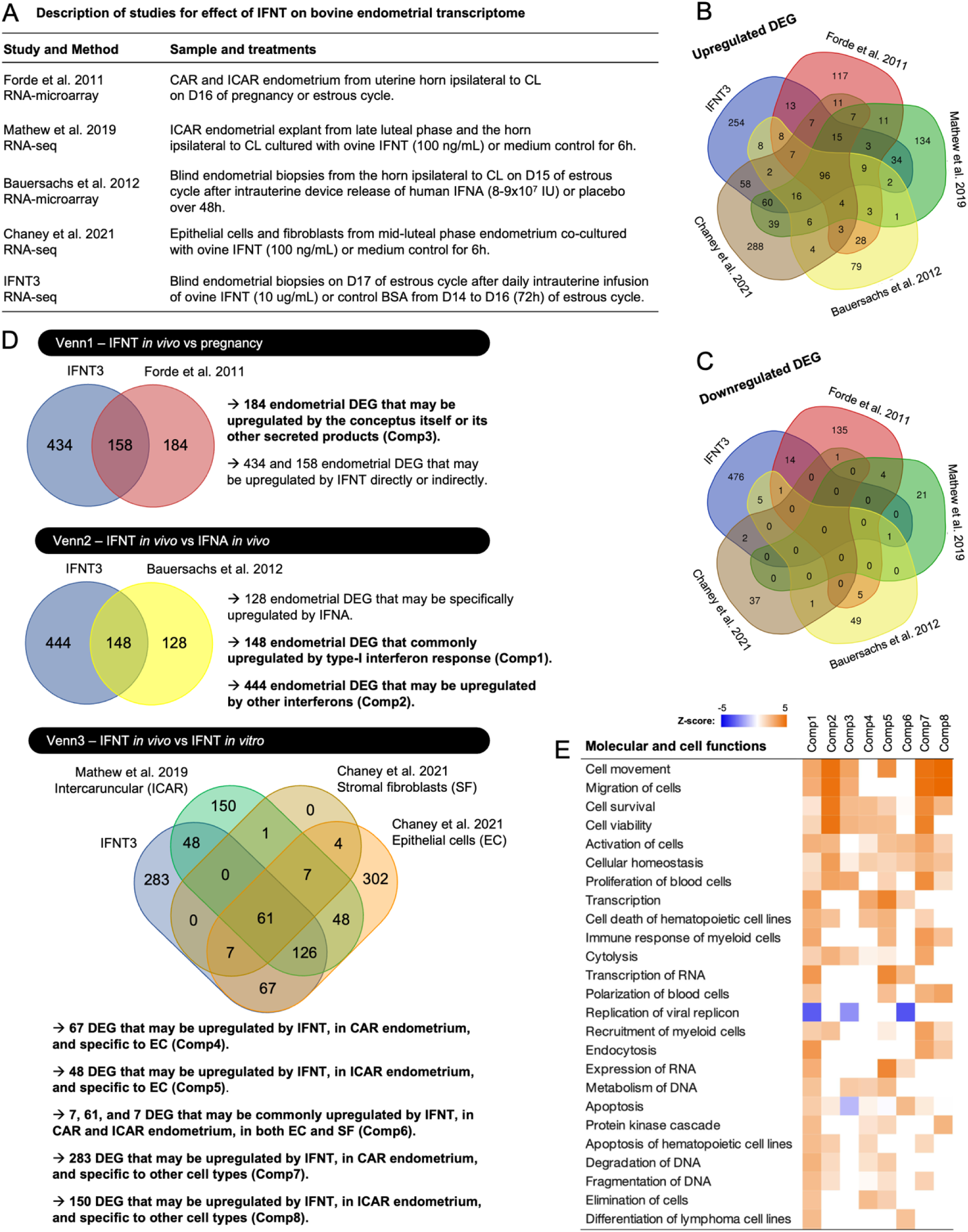
Meta-analysis of endometrial DEG in IFNT3 and DEG from experimentally similar studies. (A) Description of studies selected for comparison of the effect of IFNT on bovine endometrial transcriptome. (B) Venn diagram analysis of upregulated DEG across selected studies. (C) Venn diagram analysis of downregulated DEG across selected studies. (D) Venn diagram analysis to identify DEG regulated by the conceptus (Venn1), IFNT or other interferons (Venn2), IFNT in caruncular (CAR) and intercaruncular (ICAR) regions and in epithelial cells (EC), stromal fibroblasts (SF) and other cell types of the endometrium (Venn3). (E) Molecular and cell functions of DEGs related to type I interferon signaling (Comp1), other types of interferon signaling (Comp2), conceptus presence (Comp3), IFNT signaling in CAR EC (Comp4), IFNT signaling in ICAR EC (Comp5), IFNT signaling in CAR and ICAR SF (Comp6), IFNT signaling in CAR other cell types (Comp7), and IFNT signaling in ICAR other cell types (Comp8).

### Endometrial DEG in IFNT6 vs IFNT3

A total of 42 DEG were identified in IFNT6 compared with IFNT3 (Log2 FC FC ≥ |1| and Qvalue < 0.15). There were 16 upregulated DEG consisting of 13 characterized transcripts, 1 predicted transcript (LOC), 1 uncharacterized transcript, and 1 transcript homologous to chromosome regions in the bovine genome. The 26 downregulated DEG consisted of 22 characterized transcripts and 4 predicted transcripts (LOC). The only DEG not identified in the INTERFEROME database were ULBP11 and ULBP27.

Results also showed that prolonged IFNT signaling changed the direction of gene expression of 12 DEG that were downregulated in IFNT3 vs BSA but upregulated in IFNT6 vs IFNT3. A total of 25 DEG that include 21 upregulated and 4 downregulated DEG were identified in IFNT6 vs IFNT3 but not differentially expressed in IFNT3 vs BSA. Additionally, 5 DEG that were upregulated in IFNT3 vs BSA became downregulated in IFNT6 vs IFNT3. The top abundant genes of each group are shown in Figure 4A. In general, most DEG in this comparison did not have high TPM values, and their expression varied somewhat across replicates of IFNT6. For example, animal #3134 showed greater TPM values than other replicates in this treatment group. Nonetheless, there was no evidence for animal health or RNA sample quality issue to justify removing this animal from the transcriptome analysis. Despite the low number of DEG, IPA analysis identified progesterone (z-score = -1.9) and tretinoin (z-score = -0.5) as upstream regulators predicted to be inhibited in IFNT6. IPA identified GJA1, LRP2, SLC31A2, and ADCYAP1R1 as genes indirectly regulated by progesterone and ALDH1A3, AP1S2, GJA1, GJB5, LRAT, LRP2, TDGF1, and ADCYAP1R1 as genes indirectly regulated by tretinoin also known as retinoic acid (RA) (Figure 4B).

**Figure 4:**
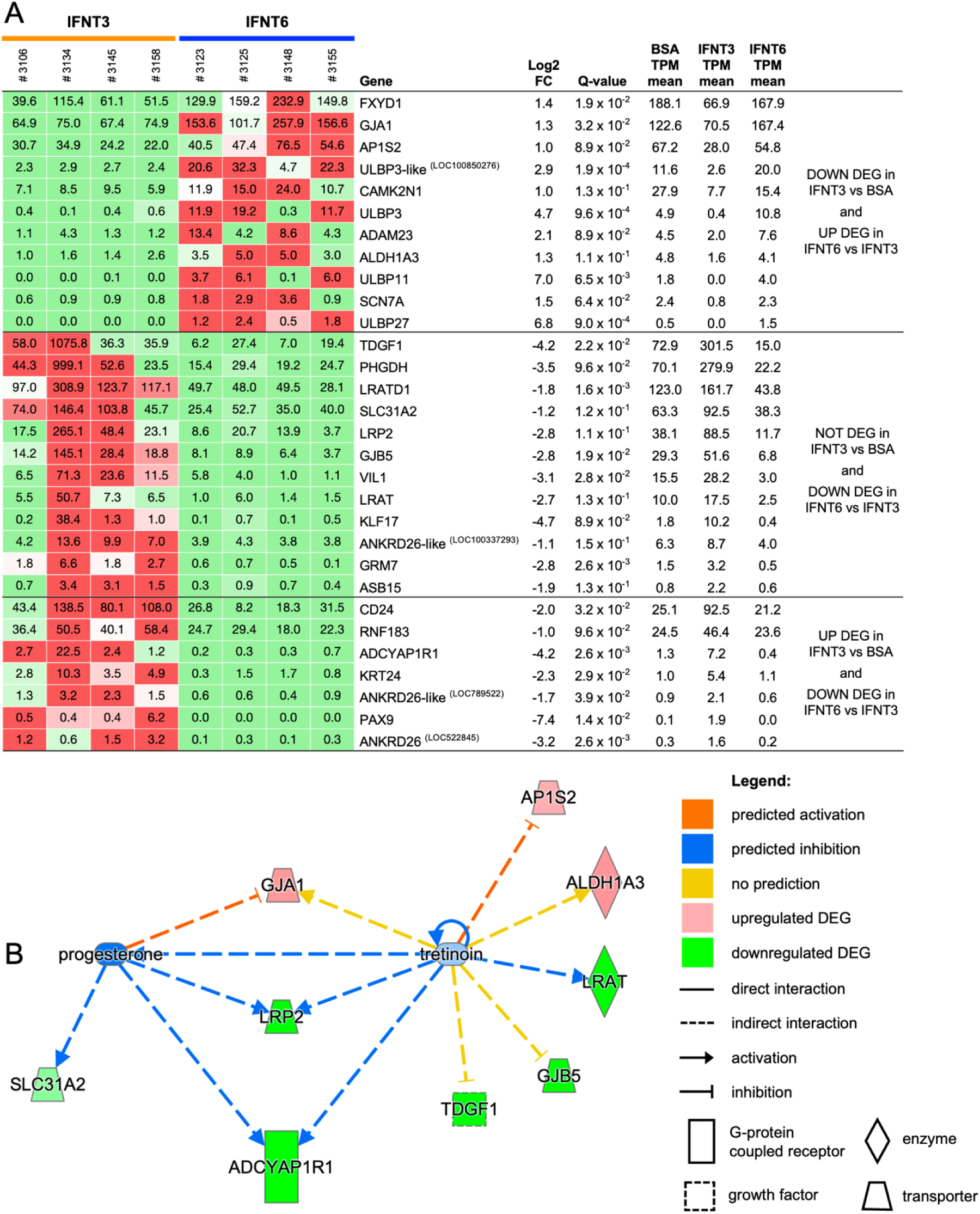
Differentially expressed genes and upstream regulators identified in endometrium of IFNT6 vs IFNT3. (A) DEG upregulated and downregulated with transcripts per million (TPM) values in each animal replicate (#) shown inside the rectangles. (B) Progesterone and retinoic acid (tretinoin) as upstream regulators of transcription predicted to be inhibited in response to IFNT6 treatment.

### Endometrial DEG in IFNT+PAG vs IFNT6

Only 4 upregulated DEG were identified in endometrium of IFNT+PAG compared with IFNT6 (Figure 5A). These DEG were did not have high TPM values, with the exception of animal #3152. There was no evidence from our records of animal health or RNA sample quality to justify removing this animal from the transcriptome analysis. Moreover, when performing DESeq2 without animal #3152, only WFDC18 was identified as a DEG. Validation of DEG via RT-qPCR using all 6 samples from treated animals showed a tendency for greater mRNA of WFDC18 and PI3 in IFNT+PAG compared to IFNT6 but no difference in expression of MUC5AC (Figure 5B-D). However, expression of these genes did not differ in endometrium from D17C, D17P, and D20P (Figure 5E-G).

**Figure 5:**
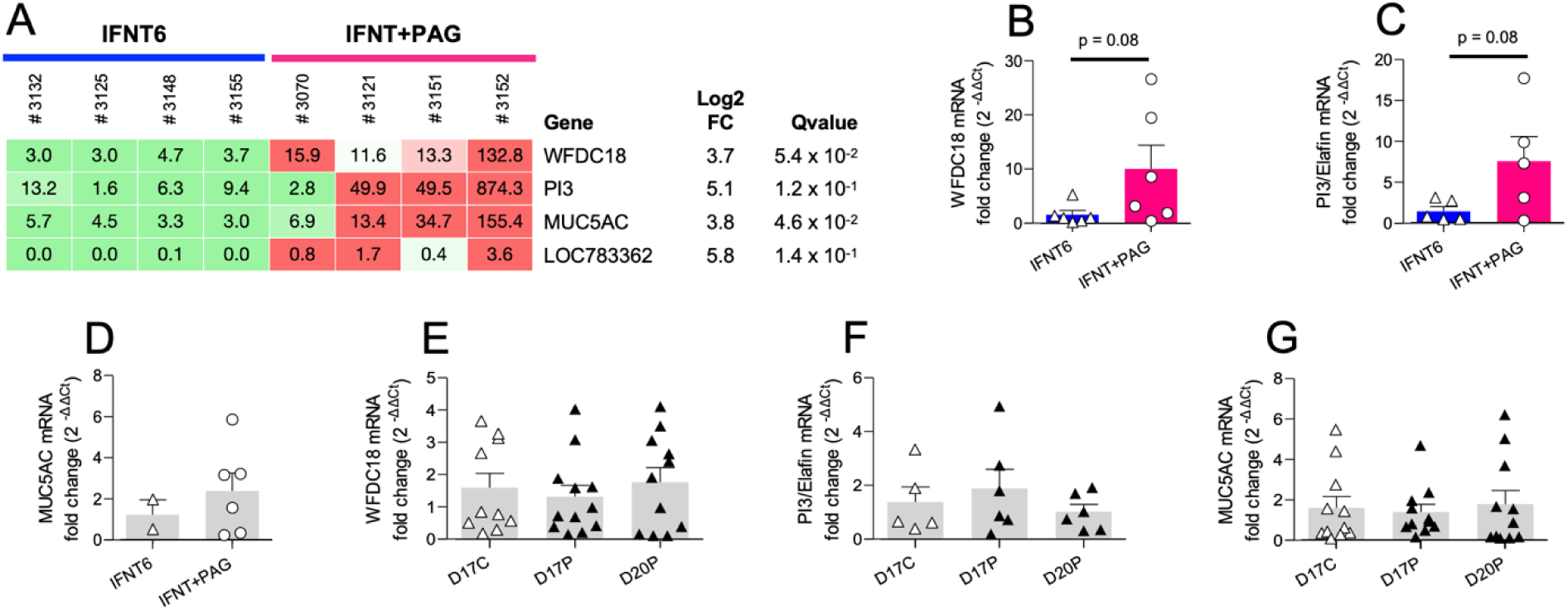
Differentially expressed genes identified in endometrium of IFNT+PAG vs IFNT6 and validated via RT-qPCR. (A) DEG with transcripts per million (TPM) values for each gene in each animal replicate (#) shown inside the rectangles. Relative gene expression of (B) WFDC18, (C) PI3, and (D) MUC5AC via RT-qPCR with endometrium samples of all animals in IFNT6 and IFNT+PAG (N=6/treatment). Relative gene expression of (E) WFDC18, (F) PI3, and (G) MUC5AC via RT-qPCR with endometrium samples of day 17 cyclic (D17C), day 17 pregnant (D17P), and day 20 pregnant (D20P) in heifers (N=12/group).

### PBL DEG in IFNT+PAG vs IFNT6

A total of 222 DEG were identified in PBL of IFNT+PAG compared with IFNT6. The 134 upregulated DEG consisted of 105 characterized transcripts, 19 predicted transcript (LOC), 9 uncharacterized transcript, and 1 transcript homologous to chromosome region. The 88 downregulated DEG consisted of 56 characterized transcripts and 26 predicted transcripts (LOC), 5 uncharacterized transcript, and 1 transcript homologous to chromosome region. The topmost abundant DEG upregulated and downregulated are shown in Figure 6A (Log2 FC ≥ |1| and Qvalue < 0.15).

**Figure 6:**
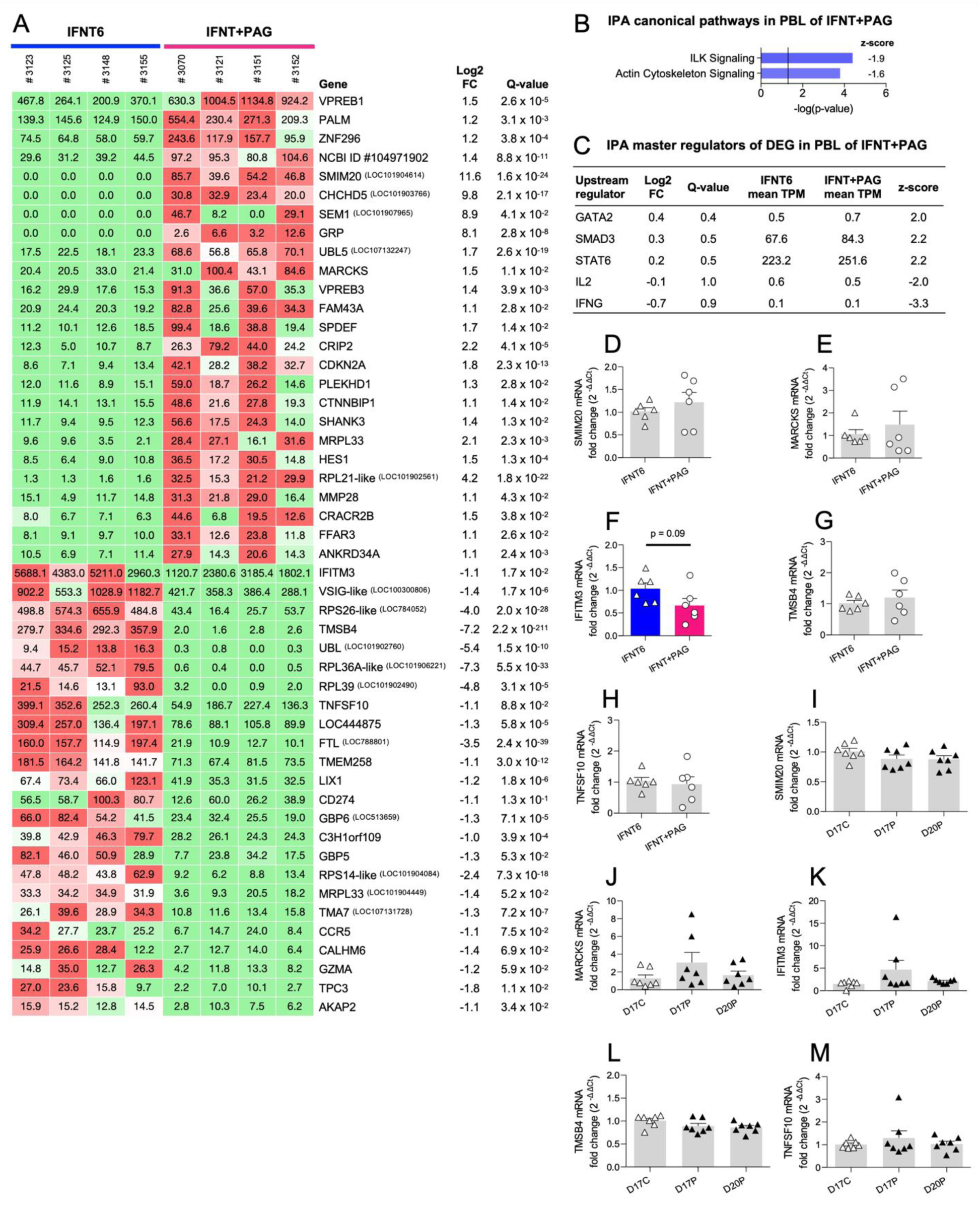
Differentially expressed genes, canonical pathways, and upstream regulators identified in blood leukocytes of IFNT+PAG vs IFNT6 and validated via RT-qPCR. (A) Most abundant DEG upregulated and downregulated in IFNT+PAG vs IFNT6, transcripts per million (TPM) values in each animal replicate (#) shown inside the rectangles. (B) IPA canonical pathways predicted to be inhibited (negative z-score, blue) by IFNT+PAG treatment with significance threshold -log (p-value of 0.05) = 1.3 (solid line). (C) IPA upstream regulators of transcription predicted to be activated in response to IFNT+PAG treatment. Relative gene expression of (D) SMIM20, (E) MARCKS, (F) IFITM3, (G) TMSB4, and (H) TNFSF10 via RT-qPCR with PBL samples of all animals in IFNT6 and IFNT+PAG (N=6/treatment). Relative gene expression of (I) SMIM20, (J) MARCKS, (K) IFITM3, (L) TMSB4, and (M) TNFSF10 via RT-qPCR with PBL samples from day 17 cyclic (D17C), day 17 pregnant (D17P), and day 20 pregnant (D20P) in heifers (N=7/group).

The IPA database recognized 108 upregulated and 54 downregulated DEG from our dataset. These DEG overlapped with 2 inhibited IPA-based canonical pathways (p-value < 0.05, z-score ≥ |1.5|) indicating that, overall, IFNT+PAG inhibited integrin linked kinase (ILK) and actin cytoskeleton signaling (Figure 6B). Upstream analysis identified 3 transcription factors that were present in the dataset and are predicted to regulate DEG in response to IFNT+PAG (Figure 6C). Validation of selected DEG via RT-qPCR using all 6 samples from treated animals showed no difference in expression, with exception of IFITM3 that showed a tendency for downregulation in response to IFNT+PAG (Figure 6D-H). However, validation of selected DEG via RT-qPCR using PBL samples from D17C, D17P, and D20P showed no difference in expression of these genes across groups (Figure 6I-M).

## DISCUSSION

This study aimed to characterize the transcription profiles of the endometrium and circulating leukocytes after treatment with IFNT and PAG. Our results indicate that short-term IFNT signaling in the endometrium (IFNT3 vs BSA) activated numerous pathways involved in interferon response, immune activation, inflammation, and cell death while downregulated genes related to vesicle transport and ECM remodeling. We also identified DEG that were not detected in previous studies and possible upstream regulators of transcription in response to IFNT. Prolonged IFNT signaling in the endometrium (IFNT6 vs IFNT3) altered the expression of genes related to cell invasion, retinoic acid (RA) signaling, and embryo implantation. Treatment with IFNT+PAG altered the expression of 4 endometrial genes that were undetectable via RT-qPCR in pregnant endometrium. On the other hand, IFNT+PAG induced numerous DEG in blood leukocytes possibly involved in inhibition of leukocyte migration, B cell differentiation and immunotolerance.

### The mechanism of IFNT signaling in bovine endometrium

As expected, all heifers treated with IFNT maintained elevated concentration of P4 in plasma confirming luteal maintenance. Initial findings suggested that IFNT preserves the CL of bovine pregnancy by suppressing endometrial transcription of OXTR (Telgmann et al., 2003), with no alteration in the expression of ESR1 (Robinson et al., 1999). However, consistent with observations in ewes (Spencer et al., 1995), analysis of the IFNT3 vs BSA dataset revealed a downregulation of OXTR (Log2 FC = -1.8; Q-value = 0.01) while indicating a tendency towards the downregulation of ESR1 (Log2 FC = -0.7; Q-value = 0.1).

In ewes, P4 and IFNT co-stimulate expression of progestamedins (Bazer et al., 2008, 2015; Bazer & Thatcher, 2017). However, endometrial expression of FGF7, FGF10, HGF, IGF1, IGF2 (stromally produced progestamedins) and their receptors did not change in response to IFNT3. Similarly, Forde et al. (2011) did not detect differences in expression of progestamedins between cyclic and pregnant cattle prior day 16. Thus, these progestamedins are unlikely to be stimulated by P4 and IFNT in the bovine endometrium during this stage of pregnancy. Hydroxysteroid 11-beta dehydrogenase 1 (HSD11B1) is stimulated in endometrium of pregnant ewes and cows (Majewska et al., 2012; Simmons et al., 2010) but was downregulated (Log2 FC = -1.2; Q-value < 0.05) in heifers treated with IFNT3 compared with control. HSD11B1 converts inactive cortisone into the active cortisol that stimulates secretion of prostaglandin E2 (PGE2) by the cow endometrium (Majewska et al., 2012). It is possible that opposite regulation of HSD11B1 mRNA between heifers in this study and prior work using cows is due to higher cortisol secretion in multiparous than primiparous cattle (Burnett et al., 2015).

### DEG specific to IFNT signaling in bovine endometrium

The INTERFEROME database contains interferon regulated genes (IRG) from experiments where different cell types/tissues from humans and mice were treated with IFNA, interferon beta (IFNB), gamma (IFNG), or lambda (IFNL) and compared with control samples. IFNT is a ruminant-specific gene with amino acid sequency identity ∼30%, ∼50%, and ∼70% similar with IFNB, IFNA, and interferon omega (IFNW), respectively (Leaman et al., 1992; Roberts et al., 1998). Thus, the five DEG in IFNT3 not identified in the INTERFEROME database may be novel IRG specific to the bovine genome and IFNT signaling.

Major histocompatibility complex class I JSP.1 (JSP.1) is one of the 28 transcribed sequences of bovine leukocyte antigen (BoLA-I) class I genes (Takeshima & Aida, 2006) with mRNA sequence matching ∼80% of nucleotides from human leukocyte antigen class I, A (HLA-A), B (HLA-B), and C (HLA-C) (NCBI Nucleotide Blast Query). These HLA molecules are present in the INTERFEROME database as IRG. During pregnancy, HLA-G isoform engages with natural killer (NK) cell receptors at the implantation sites of human and mouse and induces secretion of angiogenic and tissue growth factors (Fu et al., 2017; Bouteiller, 2015). In cattle, NK cells are upregulated during pregnancy and represent ∼50% of uterine CD45^+^ leukocytes (Vasudevan et al., 2017). Upregulation of JSP.1 mRNA was also observed in endometrial cells exposed to IFNT both *in vitro* and during early pregnancy (Wang et al., 2021; Zhu et al., 2017). These findings suggest that JSP.1 is an ISG that may be involved in regulation of NK cell function and embryo implantation.

Lymphocyte antigen 6 family member L (LY6L), was also upregulated in endometrial cells cultured with IFNT (Chaney et al., 2021). Expression pattern and cellular function of LY6L is unknown, however, other proteins in the LY6 gene family are upregulated in different types of cancer, are expressed in leukocytes, and have immunoregulatory roles (Loughner et al., 2016; Upadhyay, 2019). The X-C motif chemokine ligand 2 (XCL2) is a chemokine constitutively expressed and upregulated upon activation of NK cells, CD4, and CD8 lymphocytes (Hennemann et al., 1999; Kelner et al., 1994). XCL2 upregulation is found in different cancers and is positively associated with cell infiltration by promoting immune cell proliferation and invasion (Cao et al., 2023). The receptor for XCL2, XCR1, is expressed in dendritic cells (Lei & Takahama, 2012) suggesting crosstalk in NK cell activation and dendritic cell recruitment (Kroczek & Henn, 2012). Both NK and dendritic cells (MHC-II^+^) were abundantly upregulated in the endometrium of pregnant cattle (Kamat et al., 2016; Vasudevan et al., 2017). Thus, XCL2 may be a novel IFNT-stimulated gene mediating NK-dendritic cell communication in early pregnancy.

Little is known about coiled-coil domain containing 188 (CCDC188), but its homozygous loss of function was found in individuals with retinitis pigmentosa (Yi et al., 2020). This disease is characterized by progressive degeneration of rod photoreceptors possibly via activation of apoptosis (Newton & Megaw, 2020). Therefore, CCDC188 downregulation might participate in interferon-induced cell death signaling (Chawla-Sarkar et al., 2003) via apoptosis. Downregulation of epidermal growth factor like domain multiple 8 (EGFL8) corelated with greater metastasis in cancer and its expression is negatively associated with Notch signaling (Subhan et al., 2020; Wu et al., 2021) that regulates cell proliferation, fate, and death (Moldovan et al., 2021). Evidence suggests that IFNG might inhibit proliferation of hematopoietic cells by interfering with activation of Notch signaling (Qin et al., 2019). Thus, EGFL8 downregulation might be involved in cell differentiation of endometrium during bovine pregnancy.

### Abundant DEG downregulated by IFNT

The top upregulated genes in response to IFNT3 correspond to classical ISGs (such as ISG15, RSAD2, USP18, and genes from MX, OAS, IFIT, and IRF families). Interferons also inhibit gene expression, but interferon down-regulated genes (IDG) are less characterized (Schoggins, 2019). Over 500 DEG were repressed in response to IFNT3 while up to 160 DEG were downregulated in experimentally similar studies. Ephrin B3 (EFNB3) was the only downregulated gene in our dataset reported previously (Chaney et al., 2021). Despite having a Log2 FC = -2.4, EFNB3 mRNA was not abundant. Little is known about EFNB3, but according to the Human Protein Atlas, it is expressed across different cell types of the endometrium (Karlsson et al., 2021) and is proposed to facilitate lung cancer invasion (Efazat et al., 2016) and prostate tumor progression (Kakarla et al., 2022).

Most IDG in our dataset were lowly abundant, with the exception of myosin heavy chain 11 (MYH11), deleted in malignant brain tumors 1 (DMBT1), and peptidase inhibitor 3-like (PI3-like). MYH11 is a contractile protein, expressed abundantly in smooth muscle and less in stromal cells of the human endometrium (Karlsson et al., 2021). MYH11 was downregulated in receptive (gestational day 15) compared to pre-receptive (gestational day 5) endometrium of dairy goats (Zhang et al., 2015). It is possible that downregulation of MYH11 by IFNT signaling reinforces uterine quiescence driven by P4 (Soloff et al., 2011). DMBT1 is downregulated in endometrial cancer (Jacob et al., 2021) and involved in suppression of cervical and ovarian cancer (Ma & Zhao, 2020; Zhang, 2019) as well as protection against microbial invasion in mucosal tissue (Madsen et al., 2010). DMBT1 mRNA was stimulated by estrogen treatment of ovariectomized monkeys and rats (Tynan et al., 2005) while DMBT1 protein expression in the oviducts was greater during the porcine follicular phase (Ambruosi et al., 2013). Interestingly, DMBT1 transcripts were greater in heifers compared to lactating cows on D19 of pregnancy (Bauersachs et al., 2017). Finally, PI3 (also known as Elafin) is expressed abundantly in stromal neutrophils (King et al., 2003) and luminal and glandular epithelial cells (Uhlén et al., 2015). On day 7 of the estrous cycle, non-lactating beef cows with a large corpus luteum showed downregulation of PI3 compared with animals with a small corpus luteum (Mesquita et al., 2014) suggesting that PI3 may be downregulated by both P4 and IFNT.

### Pathways downregulated by IFNT

Canonical pathways enriched by IRG are commonly reported to be involved in antiviral response, inflammation, immune activation, and cell death (Barber, 2000; Borden et al., 2007; Crow & Ronnblom, 2019; Lazear et al., 2019). However, IPA analysis also suggested inhibition of pathways for intrinsic prothrombin activation and GP6 signaling. Most of the genes associated to these pathways are involved in extracellular matrix (ECM) remodeling such as collagen (COL4A6, COL2A1, COL4A5, COL26A1, COL11A1, COL3A1, COL5A3, COL5A1, COL9A3, COL21A1, COL27A1), laminin (EGFLAM, LAMB1, LAMB2), and Kallikrein Related Peptidases (KLKs; KLK5, KLK4). An overlap between GP6 signaling and ECM remodeling was previously observed in endometrial cancer progression (Yadav et al., 2020). Similarly, kallikreins (KLK) are proteolytic enzymes that can cleave adhesion molecules and extracellular matrix components in cancer (Srinivasan et al., 2022). Moreover, interferons were reported to inhibit transcription of collagen (Granstein et al., 1990) and bovine endometrial cells treated with IFNT in culture showed fewer collagen fibers secreted in medium along with decreased expression of metalloprotease (MMP) 2 and 9 (Rahman et al., 2020). In our dataset, MMP-12 was the only protein downregulated in IFNT3 (Log2 FC = -4.1; Q-value = 0.03). Interestingly, evidence suggests that MMP-12 can cleave IFNG and inhibit its ability to activate receptor signaling (Chopra et al., 2019).

Other downregulated genes in our dataset relate to inhibition of pathways for calcium and soluble N-ethylmaleimide-sensitive factor attachment protein receptor (SNARE) signaling. Calcium influx mediates cell proliferation, secretion, transcription, and contraction (Bootman & Bultynck, 2020), all processes regulated in the pregnant uterus. Our results showed downregulation of calcium channels (KCNMA1, CACNA1G, CACNA2D1, CACNA1A), ATPase pumps (ATP2B2, ATP2B3, ATP2B4, ATP2B4), and other calcium related molecules (CABP1, CAMK2N1, SMOC2) in the IFNT3 treatment, suggesting that IFNT alone or IFNT+P4 affects calcium signaling. Moreover, calcium mediates vesicle fusion via the SNARE complex in neural and non-neural cells (Dingjan et al., 2022; Stojilkovic, 2005). Many genes for SNARE proteins (STX1B, SNAP25, SYT3, ESYT1, VAMP3) were downregulated in IFNT3-treated heifers. SNARE proteins also participate in cytokine secretion, degranulation, and phagocytosis of immune cells (Phatarpekar & Billadeau, 2020; Stow et al., 2006). Contrary to this finding, an increase in degranulation was observed in uterine leukocytes of pregnant heifers during early pregnancy (Vasudevan et al., 2017). Overall, our results suggest that short-term IFNT may inhibit cellular events that involve ECM remodeling, calcium transport, and SNARE proteins such as immune cell migration, uterine quiescence, and regulation of immune function during pregnancy.

### Main regulators of IFNT-induced transcription

As expected, IPA identified multiple interferon regulatory factors (IRF) and signal transducer and activator of transcription (STAT) molecules as upstream regulators of DEG in our study (Mogensen, 2019). Among IRF, IRF7 was the most upregulated gene in our dataset and is a master regulator and amplifier of type-I interferon response (Honda et al., 2005; Marie, 1998). Interferon gamma inducible protein 16 (IFI16) and PML nuclear body scaffold (PML) are known ISG (Schoggins, 2019) also identified as upstream regulators in IFNT3-treated heifers. IFI16 binds to cytosolic DNA from damaged cells and pathogens to induce STING-TBK1-IRF3 interferon response (Dunphy et al., 2018; Ka et al., 2021; Paludan & Bowie, 2013) while PML (also named TRIM19) is the main organizer of nuclear bodies that regulate a variety of cellular processes such as apoptosis, proteolysis, and gene expression (Lallemand-Breitenbach & de Thé, 2018; van Damme et al., 2010). PML was also shown to participate in expression of IFNG target genes (El Bougrini et al., 2011) and regulation of apoptosis and calcium release from endoplasmic reticulum (Giorgi et al., 2010). Both IFI16 and PML are DEG in other transcriptome studies of the bovine endometrium under IFNT treatment (Bauersachs et al., 2012; Chaney et al., 2021; Forde et al., 2022; Mathew et al., 2019).

Additionally, analysis for causal networks predicted that IFNT3 may restrict nuclear receptor corepressor 1 (NCOR1) signaling leading to inhibition of early pregnancy loss. NCOR1 repress transcription of nuclear receptors such as ESR1 (Kim et al., 2021), RAR (Horlein et al., 1995), PPARG (Battaglia et al., 2010) and IRF7 (Ahad et al., 2020) by recruiting and activating histone deacetylases (HDAC) that generate a closed conformation of the chromatin. In the bovine endometrium, NCOR1 protein expression was greater on day 6-16 of the estrous cycle and positively correlated with P4 concentration (Rekawiecki et al., 2017). NCOR1 mRNA expression has not been described in the pregnant endometrium of cattle nor was it detected in IFNT3-treated heifers. However, downregulation of REST Corepressor 2 (RCOR2) was observed in the IFNT3 group. Low expression and knockdown of RCOR2 is associated with greater inflammation (Alvarez-López et al., 2014) and formation of T regulatory cell (Xiong et al., 2020) in mice. One interpretation of these results is that IFNT-induced downregulation of this corepressor may contribute to regulation of inflammatory response seen in IFNT3 transcriptome.

### Abundant DEG regulated by prolonged IFNT

During an infection, the interferon response induces pathogen clearance but also needs to be downregulated to avoid tissue damage (Arimoto et al., 2018). Ongoing and elevated interferon signaling is often seen in chronic virus infection, cancer, and type-I interferonopathies such as systemic lupus erythematosus (SLE) (Lee-Kirsch, 2017; Snell et al., 2017). Therefore, it is intriguing that the prolonged secretion of IFNT, for about 30 days total (Bazer & Thatcher, 2017), is a normal aspect of bovine pregnancy. When comparing IFNT6 vs IFNT3, there were no differences in transcription of classic ISG (Schoggins, 2019) and negative regulators of type-I interferon signaling (Arimoto et al., 2018), suggesting continuous activation of type-I interferon signaling in IFNT6. However, the identification of 42 DEG indicate that endometrial gene expression differs after prolonged exposure to IFNT and/or P4 or due to other, indirect, actions or factors.

Around day 20 of pregnancy, trophoblast cells begin to develop a closer association with the endometrium by attaching, migrating, and fusing with luminal cells to form the syncytial epithelium of placentomes (Wooding, 1992). In sheep, luminal cells undergo apoptosis and generate gaps that are filled by multinucleated trophoblast cells (Seo et al., 2019). In cows, this syncytial epithelium is temporary and replaced by dividing luminal cells (Wooding, 1992). Multinucleated trophoblast cells continue to migrate and fuse with isolated luminal cells that quickly die (Wooding, 1992). Our hypothesis is that chronic IFNT signaling regulates genes involved with the onset of migration/attachment/fusion events. Among the most abundantly transcribed DEG in IFNT6 are CD24, ring finger protein 183 (RNF183), gap junction protein alpha 1 (GJA1), FXYD domain containing ion transport regulator 1 (FXYD1), solute carrier family 31 member 2 (SLC31A2), and LRAT domain containing 1 (LRATD1).

CD24 is a cell surface glycoprotein expressed in immune, epithelial, muscle, neural, pancreas and cancer cells (Fang et al., 2010). Evidence suggests that CD24 controls homeostatic proliferation of T cells (Liu & Zheng, 2007), it is highly expressed in B regulatory cells (Breg) (Blair et al., 2010), but its deficiency increases T regulatory cells (Treg) immunosuppressive abilities (Shi et al., 2022). CD24 is highly expressed in cancer, correlating with cell invasion and proliferation as well as reducing phagocytosis via interaction with macrophage Siglec-10 (Altevogt et al., 2021; Barkal et al., 2020). RNF183 is an E3 ubiquitin ligase, and it is upregulated in uterine corpus endometrial carcinoma and negatively correlated with infiltration of CD4+ T cells, neutrophils, and dendritic cells (Geng et al., 2020). In inflammatory bowel disease, RNF183 is also upregulated and drives inflammation by targeting NFKB inhibitor alpha (NFKBIA) for degradation and activating NFKB signaling (Yu et al., 2016). In the human endometrium, CD24 and RNF183 are most abundantly expressed in luminal and glandular cells (Karlsson et al., 2021). Thus, downregulation of these molecules suggests inhibition of immune activation under chronic IFNT signaling.

In cattle, GJA1 (also known as CX43) is localized in caruncles and placentomes (Mansouri-Attia et al., 2009; Pfarrer et al., 2006). It is also upregulated in porcine endometrium on day 12 of pregnancy (Zeng et al., 2018). Increased expression of GJA1 is an indicator of decidualization and promotes stromal cell proliferation in rodents (Kim et al., 2022; Orlando-Mathur et al., 1996; Yu et al., 2017). FXYD1 (also known as Phospholemman or PLM) is located near the basolateral membrane and regulates the kinetic properties of Na^+^/K^+^ ATPase activity (Arystarkhova et al., 2017; Bibert et al., 2008; Crambert et al., 2002). Beyond ion transport, Na^+^/K^+^ ATPase activity participates in cell junction formation, cell adhesion, motility, and proliferation (Aperia, 2007; Contreras et al., 1999; Rajasekaran & Rajasekaran, 2003; Silva et al., 2021). Both GJA1 and FXYD1 are abundantly transcribed in stromal cells of the human endometrium (Karlsson et al., 2021), therefore, they may regulate stromal remodeling during pregnancy.

SLC31A2 (or CTR2) is an ion transporter that facilitates copper mobilization from intracellular vesicles to the cytoplasm (Wee et al., 2013). In human endometrium, SLC31A2 is abundantly transcribed in macrophages (Karlsson et al., 2021) and copper is an important metal in energy generation, phagocytosis and killing ability (Stafford et al., 2013). Studies have shown upregulation (Wagner et al., 2005; White et al., 2009) and downregulation of copper transport upon IFNG signaling (Shen et al., 2018) in different disease models. However, downregulation of endometrial SLC31A2 in the IFNT6 treatment and in day 30 pregnant endometrium of buffalo (Panigrahi et al., 2020) supports the hypothesis that regulation of macrophage phagocytosis is important during implantation and placentation. Expression of LRATD1 mRNA is greater in endothelial cells, luminal and glandular epithelial cells of human endometrium (Karlsson et al., 2021). Knowledge about this molecule is limited, but LRATD1 is commonly upregulated in colorectal cancer and may play a role in cell motility (Kobayashi et al., 2006). Overall, these DEG support our hypothesis that IFNT6 may regulate genes involved with implantation processes such as cell invasion, stromal remodeling, and immune regulation.

### Prolonged IFNT treatment and RA signaling during implantation/attachment

IPA analysis of the IFNT6 data predicted inhibition of progesterone and retinoic acid signaling. Downregulation of progesterone receptor (PGR) in the endometrial luminal and superficial glandular epithelium is common prior embryo implantation/attachment in mammals and occurs by day 13 in cyclic and pregnant cattle due to prolonged exposure to P4 (Okumu et al., 2010). High P4 concentration increases expression of retinol binding proteins (RBP) and concentration of RA in the uterine lumen of pregnant cattle (Simintiras et al., 2019; Thomas et al., 1992). For the bovine conceptus, retinoid receptor signaling participates in lipid metabolism critical during elongation (Mohan et al., 2002; Ribeiro, 2018). However, knowledge about the role of RA in the bovine endometrium is limited. In humans, RA has been reported to inhibit decidualization and that endometrial stromal cells attenuate RA signaling during normal decidualization (Brar et al., 1996; Ozaki et al., 2017). Thus, it is possible that inhibition of RA signaling in bovine endometrium may be a result of prolonged IFNT signaling to facilitate implantation.

Among the DEG in IFNT6 related with RA signaling, lecithin retinol acyltransferase (LRAT) and aldehyde dehydrogenase 1 family member A3 (ALDH1A3) are enzymes that convert retinol into retinyl esters for storage and into the bioactive RA, respectively (Ruiz et al., 1999; Sima et al., 2009). However, ALDH1A3 also participates in the metabolism of glucose and amino acids (Duan et al., 2016). Other RA-related DEG include the LDL receptor related protein 2 (LRP2; also called Megalin) and teratocarcinoma-derived growth factor 1 (TDGF1; also called Cripto-1). The gene for LRP2 encodes an endocytic receptor induced by progesterone and retinoids that is involved in the uptake of lipids, cholesterol, hormones, minerals, and vitamins including RA (Liu et al., 1998; Marzolo & Farfán, 2011). LRP2 is expressed in CAR and ICAR on day 20 of bovine pregnancy (Mansouri-Attia et al., 2009; Pfarrer et al., 2006). Interestingly, the DEG adaptor related protein complex 1 subunit sigma 2 (AP1S2) involved in clathrin-coated vesicle formation was also found to mediate apical protein localization of LRP2 (Gravotta et al., 2019). TDGF1 is an epidermal growth factor and co-receptor in transforming growth factor beta (TGFB) signaling also important for early embryogenesis, epithelial-mesenchymal transition, cell migration, and angiogenesis (Klauzinska et al., 2014). Mice lacking TDGF1 show impaired decidualization, implantation and placental vasculature (Shafiei & Dufort, 2021). Interestingly, a positive correlation was detected between expression of TDGF1 and level of invasiveness of trophoblast tissue in ampullary pregnancies (Cabar et al., 2022). Overall, the literature supports that the DEG in IFNT6 may facilitate cellular processes required for decidualization and implantation that are equivalent to events for embryo attachment in bovine pregnancy.

### Limited response to PAG in the endometrial transcriptome

A few studies have reported a function for PAG in regulating gene expression of ruminant endometrium. However, due to experimental design, it is unclear if the results obtained in these studies are physiologically relevant or directly caused by PAG. For example, Vecchio et al. (Del Vecchio et al., 1990) reported that bovine endometrium from day 16 of estrous cycle cultured with a mixture of first trimester placental PAG (PSPB), without exposure to IFNT, tended to show elevated PGF2A and greater PGE2 secretion compared to controls. Later, Weems et al (2003) treated caruncles and placentomes from pregnant ewes *ex vivo* (exposed in utero to IFNT) with PAG, and reported no effect of PAG on PGF2A and PGE2 between day 13 and 50 of pregnancy. Thus, PAG seemed not to alter endometrial secretion of prostaglandins during early pregnancy. Another study reported that porcine PAG-2 and PAG-12 showed proteolytic activity *in vitro* under acidic pH, but without the presence of any conceptus secretions (Telugu et al., 2010). Regarding effects of PAG on endometrial gene expression, Wallace et al. (2019) treated day 18 endometrium from non-pregnant and pregnant heifers with PAG and observed increased expression of genes involved with ECM remodeling. However, pregnant endometria were only exposed to IFNT in utero and not during culture and treatment with PAG *ex vivo*. Moreover, most of the differences in gene expression induced by PAG treatment were only detected at 96 hours of culture when expression of ISG were at their lowest (Wallace et al., 2019). Our transcriptome analysis and the work of Rahman et al. (2020) showed that IFNT downregulated expression of metalloproteases and begs the question of whether results observed *in vitro* by Wallace et al. (2019) would apply to physiological pregnancies when endometrium is concomitantly exposed to PAG and IFNT and in the presence of physiological concentrations of progesterone.

The transcriptome analysis of Mansouri-Attia et al. (Mansouri-Attia et al., 2009) comparing endometrium of day 20 of pregnancy with day 20 of estrous cycle identified numerous DEG in response to pregnancy. However, the control samples from day 20 estrous cycle were not exposed to IFNT and animals likely had low progesterone and elevated estradiol concentration in plasma compared to samples from pregnant animals. These physiological differences possibly affected the transcriptome profile obtained in that study, making it harder to determine the role of PAG in the endometrium of pregnancy. Our transcriptome analysis mimicked the physiological timing and concentration of P4, IFNT, and PAG secretion in the uterus. However, results suggest that PAG have minimal effects on gene expression in bovine endometrium. Wooding (Wooding, 1984) described that invading trophoblast cells in ewes appear to secrete PAG near endometrial blood vessels. Thus, it is possible that PAG signaling targets circulating leukocytes (Hoeben et al., 1999) or extrauterine tissues such as the CL (Weems et al., 1998) during pregnancy.

### Abundant DEG are regulated by PAG in circulating leukocytes

Among DEG upregulated by IFNT+PAG in circulating leukocytes, the genes for paralemmin (PALM) and V-Set Pre-B Cell Surrogate Light Chain 1 (VPREB1) had the highest number of transcripts. According to the Human Protein Atlas, PALM is specific to non-classical monocytes while VPREB1 is specific to and mainly expressed in naïve and memory B-cells (Karlsson et al., 2021). Despite limited information about the function of these genes, PALM participates in plasma membrane dynamics such as formation of filopodia (Arstikaitis et al., 2008; Kutzleb et al., 1998) while VPREB1 and VPREB3 are involved in early steps of B-cell differentiation (Lee et al., 2021; Rossi et al., 2006). Interestingly, transcripts of small integral membrane protein 20 (SMIM20) and coiled-coil-helix-coiled-coil-helix domain containing 5 (CHCHD5) were undetected in the IFNT6 treatment and they were upregulated in PBL of IFNT+PAG. SMIM20 (also known as MITRAC7) is a chaperone specific to assembly of cytochrome c oxidase (COX) in the inner mitochondrial membrane (Dennerlein et al., 2015) while CHCHD5 is an isoform of nucleus-encoded protein that is imported to the mitochondrial intermembrane space (Modjtahedi et al., 2016). The function of CHCHD5 is unknown, but other isoforms are described to participate in regulation of COX activity, mitochondrial translation, apoptosis, cell migration, and mitochondrial morphology (Modjtahedi et al., 2016).

Interferon induced transmembrane protein 3 (IFITM3) and V-set and immunoglobulin domain containing-like protein (VSIG-like) are downregulated DEG in IFNT+PAG with the highest number of transcripts. IFITM3 was upregulated in IFNT3 vs BSA (Log2 FC = 2.3; Q-value < 0.001) and abundantly transcribed yet not differentially expressed in the other treatment comparisons. Conversely, PAG treatment decreased IFITM3 expression in PBL. IFITM3 is an ISG known to have antiproliferative function and inhibit viral fusion with cells (Bailey et al., 2014). It is also suggested that IFITM proteins promote T cell differentiation and that their absence decreases migration, cytokine secretion and inflammation (Yánez et al., 2019). Similarly, knockdown of IFITM3 expression reduced cell migration and proliferation in gastric tumor cells (Hu et al., 2014). VSIG proteins are recognized as immune checkpoint receptors and their expression correlate with B cell and macrophage infiltration as well as inhibition of T cell activation (Zhou et al., 2022). Along with VSIG, CD274 (or programmed cell death 1 ligand 1; PD-L1) was also downregulated in PBL exposed to PAG. CD274 is also an immune inhibitory receptor, upregulated under interferon signaling (Garcia-Diaz et al., 2017). Another noticeable DEG is thymosin beta 4 (TMSB4) which was downregulated by ∼147 fold (Log2 FC = -7.2) in response to IFNT+PAG. TMSB4 is an actin binding protein involved in cell migration, adhesion, and differentiation of mononuclear cells in peripheral blood (Ito et al., 2009; Liao et al., 2022; Mælan et al., 2007). Actin alpha 1 (ACTA1) was also downregulated in response to IFNT+PAG treatment.

### Possible function of PAG in circulating leukocytes

Pathways analysis of the DEG in the IFNT6+PAG treatment identified inhibition of actin cytoskeleton and ILK signaling pathways. However, the DEG associated with these pathways were lowly abundant (ACTA1, MYH1-3, MYH14, MYO18B, TTN, PPP2R2C), except for TMSB4. Nonetheless, along with the abundant downregulated DEG discussed in the previous section, they would contribute to inhibition of leukocyte migration. Actin and myosin are important molecules for cell morphology and movements in leukocytes (Vicente-Manzanares & Sánchez-Madrid, 2004). Similarly, ILK signaling promotes actin filament bundling that generates contraction force and mechanical signals involved in cell adhesion (Vaynberg et al., 2018). Thus, it is possible that PAG inhibit leukocyte recruitment to the uterus during early pregnancy of cattle.

Upstream analysis identified activation of transcription factors GATA binding protein 2 (GATA2), STAT6, and SMAD family member 3 (SMAD3). GATA2 is crucial for hematopoiesis and mutations in GATA2 gene leads to myelodysplasia and deficiency in B cells, monocytes, NK cells, and dendritic cells (Calvo & Hickstein, 2023; Nováková et al., 2016; Tsai et al., 1994). Moreover, STAT6 activation is mainly induced by IL4 that, in turn, supports B cell development, activation, and tolerance (Wang et al., 2021). Thus, predicted activation of GATA2 and STAT6 along with upregulation of genes VPREB1 and VPREB3 in response to IFNT+PAG suggest that PAG may affect B cell development in peripheral blood. Similarly, Hoeben et al. (Hoeben et al., 1999) reported that PAG can inhibit growth of mononuclear bone marrow cells and peripheral blood leukocytes *in vitro*. However, Vasudevan (2016) reported no effect of IFNT and PSPB (PAG) alone or in combination on proliferation of peripheral immune cells *in vitro*. Proteins of the SMAD family are effectors of TGFB signaling important for immunotolerance and resolution of inflammation (Li et al., 2006). SMAD3 induces Treg cells and myelopoiesis (Giroux et al., 2011; Martinez et al., 2009). Emerging evidence also suggests that B regulatory cells (Breg) participate in secretion of TGFB and generation of Treg (Lee et al., 2014; Rosser & Mauri, 2015). Moreover, mRNA expression of anti-inflammatory cytokines IL10, IL4, TGFB and tolerogenic marker IDO are upregulated in peripheral blood mononuclear cells (PBMC) and neutrophils between day 10-18 of pregnant versus cyclic cows (Mohapatra et al., 2020; Yang et al., 2016). Thus, PAG may induce anti-inflammatory signals and regulatory lymphocytes to suppress immune response to the conceptus and lower peripheral inflammation.

## Conclusion

This study identified the transcription profile of the endometrium and circulating leukocytes in response to IFNT and PAG. Abundant production of PAG by the ruminant conceptus has been known for nearly 40 years, yet their function remains an enigma. Overall, our results showed that: (1) Acute IFNT signaling activated immune response pathways and may inhibit ECM remodeling and vesicle transport in the endometrium; (2) 5 genes were identified that may be specific to the bovine genome and IFNT signaling; (3) prolonged IFNT signaling shifts the expression of some genes that may to be involved in inhibition of retinoic acid signaling altered and cell invasion during implantation/attachment; (4) PAG altered gene expression in blood leukocytes more than in endometrium; and (5) PAG may stimulate leukocyte development and TGFB signaling and inhibit interferon signaling and migration in blood leukocytes. The strength of the present study was the ability to apply intrauterine treatments that mimic the physiological timing and concentration of IFNT and PAG *in vivo* pregnancies. Weaknesses include the use of PAG isolated from first trimester placenta that may not reflect of the mixture of PAG isoforms secreted in early pregnancy. It is also important to consider that the conceptus may alter gene expression of endometrial cells via mechanical stimuli or secretion of other proteins and molecules. Nonetheless, our results suggest that during early pregnancy IFNT rather than PAG is a major regulator of endometrial gene expression, but PAG can alter the transcriptome of immune cells in circulation when administered in physiological concentrations. Mechanistic research is required to explore the role that the key DEG identified in this study play in the endometrium and blood leukocytes. Overall, our results contribute to the understanding about the effects of conceptus secretions on uterine and immune function during a period of high embryo loss.

**Table 1:**
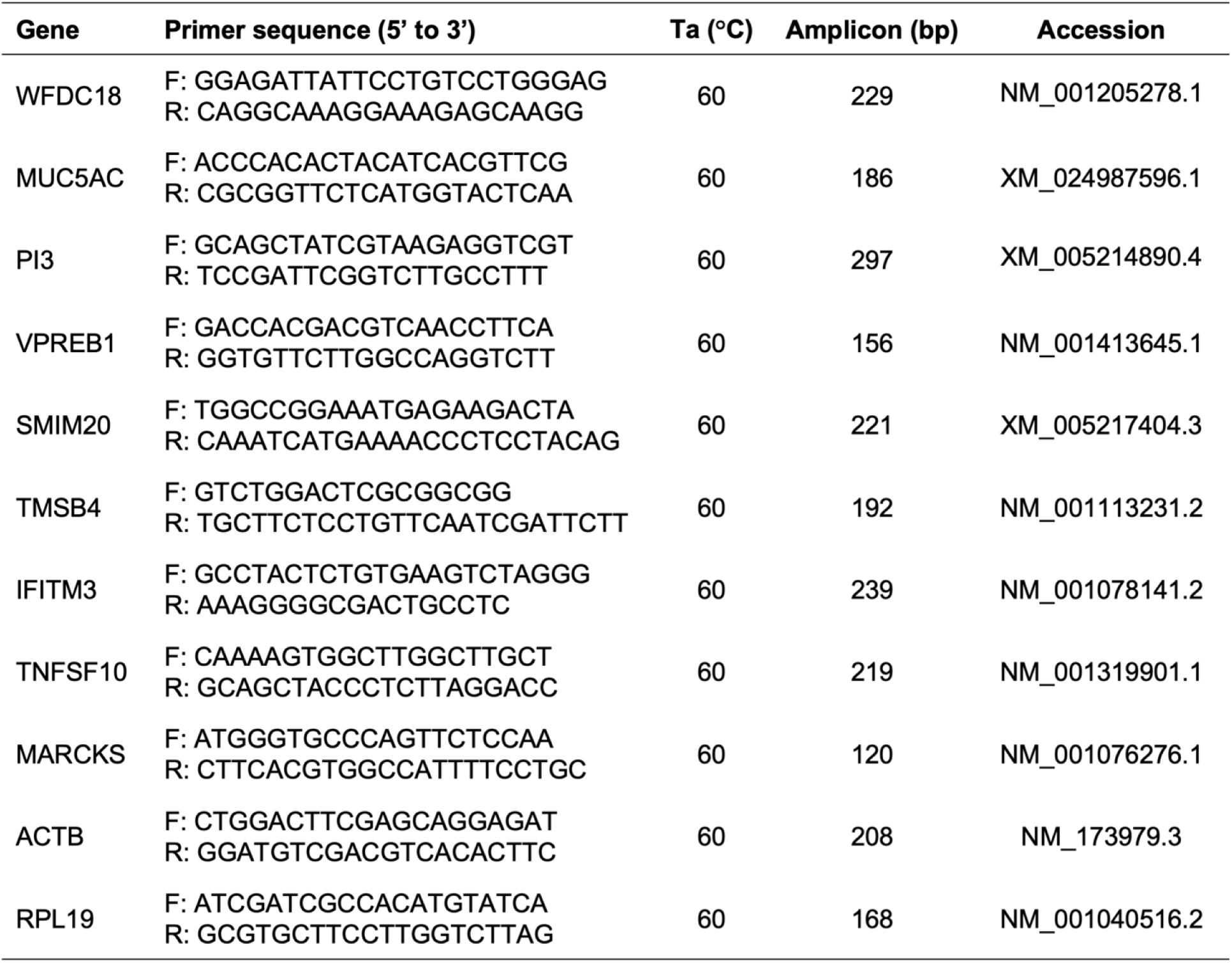
Primers used in RT-qPCR analysis.

## Acknowledgements

We thank Dr. Joy Pate for her critique of this manuscript and Nadine Houck, Travis Edwards, and Ty Montgomery for assistance in handling animals during experimental treatments and sample collection.

## Data availability

Data related to this article is incorporated into main figures and deposited in NCBI’s Gene Expression Omnibus (GEO) under the accession number GSE241886 or the link https://www.ncbi.nlm.nih.gov/geo/query/acc.cgi?acc=GSE241886. Data in GEO includes raw files, list of DEG and DEG selected for meta-analysis.

## Author contributions

TO conceived the research question and experimental design. MIS conducted the experiment, data analysis and interpretation, and the writing of the manuscript. TO provided significant intellectual contributions during manuscript content review. All authors read and approved the submitted version.

